# Inhibiting adenine synthesis attenuates glioblastoma cell stemness and temozolomide resistance

**DOI:** 10.1101/2021.06.21.449341

**Authors:** Simranjit X. Singh, Rui Yang, Kristen Roso, Landon J. Hansen, Changzheng Du, Lee H. Chen, Paula K. Greer, Christopher J. Pirozzi, Yiping He

**Author notes:** Corresponding Author: Yiping He, PhD, 203 Research Drive, Medical Science Research Building 1, Room 199A, Duke University Medical Center, Durham, NC 27710.

## Abstract

Glioblastoma (GBM) is a lethal brain cancer exhibiting high levels of drug resistance, a feature partially imparted by tumor cell stemness. Recent work shows that homozygous *MTAP* deletion, a genetic alteration occurring in about half of all GBMs, promotes stemness in GBM cells. Exploiting MTAP loss-conferred deficiency in adenine salvage, we demonstrate that transient adenine blockade via treatment with L-Alanosine (ALA), an inhibitor of *de novo* adenine synthesis, attenuates stemness of *MTAP*-deficient GBM cells. This ALA-induced reduction in stemness is accompanied by compromised mitochondrial function, highlighted by diminished spare respiratory capacity. Direct pharmacological inhibition of mitochondrial respiration recapitulates the effect of ALA on GBM cell stemness, suggesting ALA targets stemness partially via affecting mitochondrial function. Finally, in agreement with diminished stemness and compromised mitochondrial function, we show that ALA sensitizes GBM cells to temozolomide (TMZ) *in vitro* and in an orthotopic GBM model. Collectively, these results identify critical roles of adenine supply in maintaining mitochondrial function and stemness of GBM cells, highlight a critical role of mitochondrial function in sustaining GBM stemness, and implicate adenine synthesis inhibition as a complementary approach for treating *MTAP*-deleted GBMs.

## INTRODUCTION

Glioblastoma (GBM) is the most prevalent and lethal primary brain tumor [1]. GBM patients have poor overall survival, owed in part to GBM’s heterogeneous and drug resistant nature [2]. Heterogeneity and drug resistance in GBM is partly driven by the presence of brain tumor initiating cells (BTICs), also known as GBM stem-like cells (GSCs) [3–6], as evidenced by their resistance to radiation and chemotherapy [3, 5, 7]. Recent studies have provided critical insights into the genesis and maintenance of BTICs and offered promising targeting strategies, including epigenetic and metabolic approaches [8–11].

Our recent study has shown that homozygous deletion of *MTAP* (methylthioadenosine phosphorylase), a genetic alteration occurring in about 50% of GBMs, promotes the stemness of GBM cells and leads to more aggressive tumors [12]. MTAP functions in the adenine salvage pathway, and loss of this enzyme renders cells sensitive to adenine deprivation [12, 13]. In preclinical models, pharmacological inhibition of *de novo* adenine synthesis was efficacious as a monotherapy against *MTAP*-deficient GBM and reduced the percentage of GBM cells positive for CD133, a glioma stem cell marker [4, 12]. However, the role of adenine supply in maintaining stemness in *MTAP*-deficient GBMs and the underlying cellular effects of adenine synthesis inhibition remain unclear.

Here, we use patient-derived *MTAP*-deficient GBM models to show that transient low-dose *de novo* adenine synthesis inhibition via L-Alanosine (ALA) attenuates GBM stemness. We demonstrate that this effect is partially mediated by attenuated mitochondrial respiration and spare respiratory capacity. Importantly, consistent with reduced stemness and mitochondrial function, we show that ALA treatment sensitizes *MTAP*-deficient GBM cells to temozolomide (TMZ), highlighting a targeted strategy to improve standard-of-care treatment.

## RESULTS AND DISCUSSION

To investigate the role of adenine supply in maintaining stemness, we employed naturally *MTAP*-deficient patient-derived GBM cell lines (GBM 12-0160 and GBM 12-0106) [12], which were cultured in neural stem cell (NSC) medium to maintain a more glioma stem-like cell state. We subjected these cell lines to extreme limiting dilution assay (ELDA) analysis [14], seeding 33, 11, 3, and 1 cell per well, treated with 0.125 μM or 0.25 μM ALA. These doses exhibit minor effects on proliferation, removing the confounding variable of cell growth inhibition in assessing changes in stemness (**Supplementary Figure 1A**). Overall, the ELDAs demonstrate that ALA treatment leads to a significant dose-dependent reduction in stem cell frequency (**Figure 1A**). Furthermore, ALA treatments (at 0.25 μM) reduce the number of spheres formed per well for both cell lines (**Figure 1B**). Representative images of tumor spheres derived from each GBM cell line qualitatively indicate a reduction in tumor sphere size with ALA treatment (**Figure 1C**). These results demonstrate that the presence of ALA during the sphere-forming period (two to three weeks) decreases stem cell frequency and sphere formation capacity of our GBM cells.

**Figure 1.**
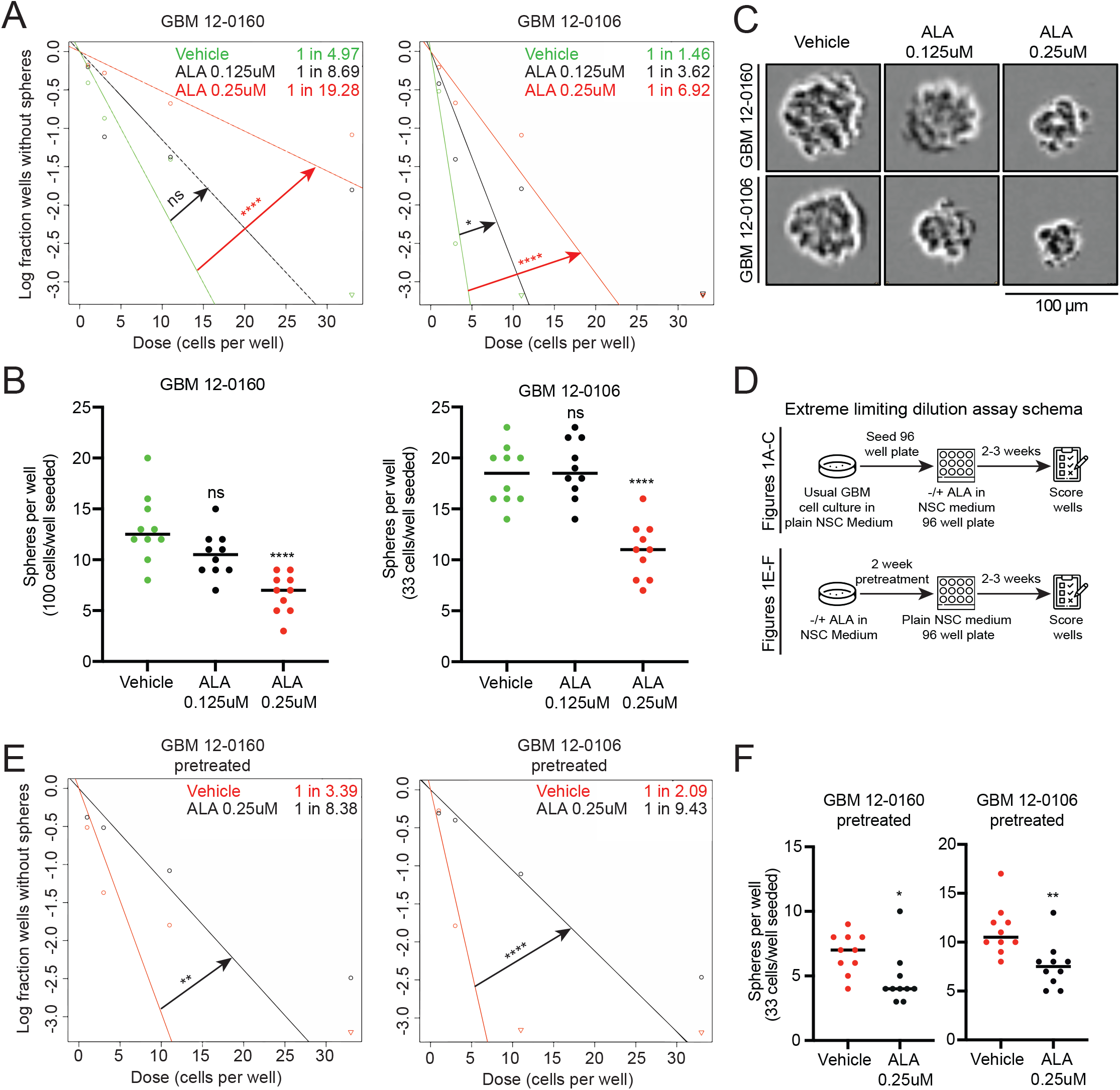
Inhibition of *de novo* adenine synthesis using L-Alanosine (ALA) decreases stem cell frequency and sphere-forming capacity of *MTAP*-deficient GBM cells. **(A)** Extreme limiting dilution assays (ELDAs) show ALA treatment decreases stem cell frequency in two *MTAP*-deficient patient-derived GBM cell lines. Stem cell frequency (1 in n) for each condition shown on plot. 33, 11, 3, 1 cells seeded per well, *n*=12 replicates per condition. **(B)** ALA treatment decreases the number of spheres formed per well in ELDA. 100 cells seeded per well (left, GBM 12–0160) or 33 cells seeded per well (right, GBM 12–0106), *n*=10 replicates per condition. **(C)** Representative images of tumor spheres derived from GBM 12–0160 and GBM 12–0106. Scale bar, 100μm. **(D)** Schema depicting design of ELDAs. Assays shown in **(A-C)** done with concurrent ALA treatment in 96 well plates; assays shown in **(E-F)** done with 2 weeks ALA pretreatment followed by seeding cells in neural stem cell (NSC) medium without ALA in 96 well plates. **(E)** Extreme limiting dilution assays (ELDAs) show 2-week pretreatment with ALA decreases stem cell frequency in GBM cells. Stem cell frequency (1 in n) for each condition shown on plot. 33, 11, 3, 1 cells seeded per well, *n*=12 replicates per condition. **(F)** 2-week pretreatment with ALA decreases the number of spheres formed per well in ELDA. 33 cells seeded per well, *n*=10 replicates per condition. Data shown are mean +/− SEM. Data analyzed using ELDA Chi-square test **(A and E)**, One-Way ANOVA followed by multiple t-tests **(B and F)**; ns: not significant, \**P* < 0.05, \*\**P* < 0.01, \*\*\*\**P* < 0.0001.

To determine whether ALA confers a transient or endured stemness reduction, we conducted ELDAs with ALA pretreatment. We pretreated GBM cells for two weeks with vehicle or with ALA (0.25 μM) before seeding the pretreated cells in NSC medium without ALA for ELDA (**Figure 1D**). Remarkably, we found that GBM cells pretreated with ALA also display a reduction in stem cell frequency and sphere forming capacity (**Figure 1E, F**). These results suggest that even transient ALA treatment confers a sustained diminishment of stemness.

To investigate the underlying cellular impacts of long term ALA treatment, we conducted mRNA-sequencing on two GBM cell lines treated with ALA (at 0.25 μM) for two weeks. Interestingly, while no differential expression was detected for a majority of genes in cells treated with ALA (**Supplementary Table 1**), ALA treatment results in an approximately 20% reduction in transcriptional output from mitochondrial DNA (mtDNA), in comparison to output from the other chromosomes (**Figure 2A**). PCR quantification of the mtDNA showed that mtDNA copy number is unaffected by ALA treatment (**Supplementary Figure 1B**). To test whether reduced mtDNA transcriptional output is indicative of mitochondrial function deficiency, we utilized Seahorse XF analysis to conduct a mitochondrial stress test on the GBM cells treated with ALA. Initially, we sought to test the effects of acute 24-hour ALA treatment, and found that high doses of ALA (3 μM and 10 μM) moderately reduce maximal oxygen consumption rate (OCR) (**Supplementary Figure 2A**). Furthermore, the addition of exogenous adenine to cells undergoing ALA treatment rescues this reduction in maximal OCR, suggesting this reduced OCR is due to adenine shortage, instead of off-target effects of ALA (**Supplementary Figure 2B)**. Next, we aimed to assess the impact of transient 2-week ALA pretreatment on OCR. Notably, we found that pretreatment of GBM cells with low doses of ALA (0.25 μM and 0.5 μM) results in significantly attenuated maximal OCR (**Figure 2B**), and eliminates mitochondrial spare respiratory capacity (**Figure 2C**). In this experiment, OCR was measured in media free of ALA; therefore, these results suggest a lasting effect of adenine shortage on mitochondrial function in GBM cells, reminiscent of the effects on GBM stemness.

**Figure 2.**
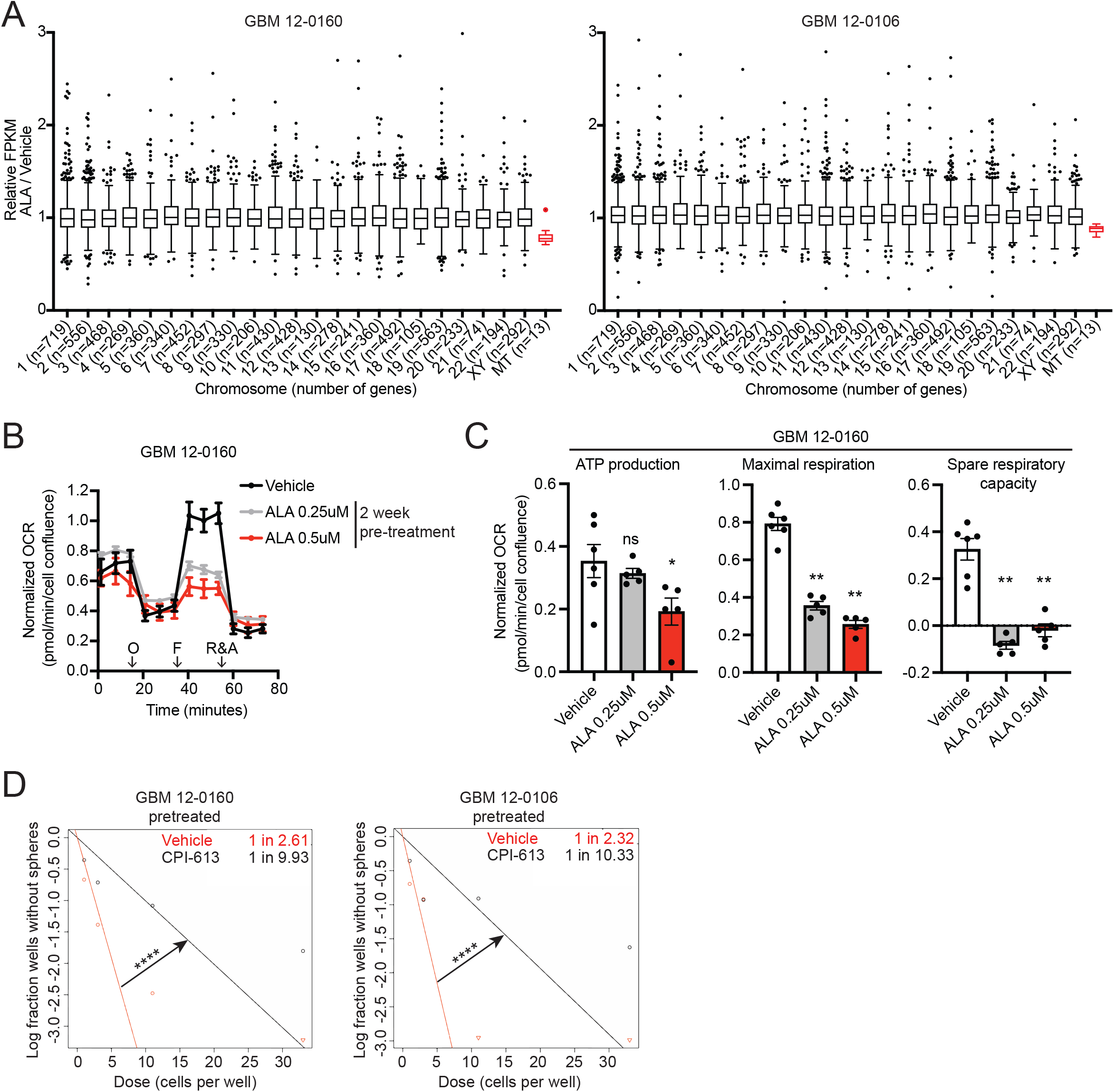
Inhibition of *de novo* adenine synthesis using L-Alanosine (ALA) reduces GBM cells’ mitochondrial respiration and eliminates spare respiratory capacity. **(A)** mRNA-seq analysis of patient-derived GBM cells pretreated for 2 weeks with ALA 0.25 uM shows that ALA decreases expression of mitochondrial DNA-encoded (MT) genes. Genes with FPKM of at least 10 were included in analysis. Outlier genes (*n*=3 genes) with relative FPKM>3 excluded from plot. *P* < 0.0001 for MT vs each other chromosome. **(B)** Seahorse XF analysis of GBM 12–0160 pretreated for 2 weeks with ALA 0.25 uM or 0.5 uM shows ALA reduces maximal respiration. *n*=6 replicates for Vehicle, *n*=5 replicates for each other condition. O: Oligomycin, F: FCCP, R&A: Rotenone and Antimycin A. **(C)** Two-week pretreatment with ALA reduces ATP production, maximal respiration, and spare respiratory capacity in GBM 12–0160. *n*=6 replicates for Vehicle, *n*=5 replicates for each other condition. **(D)** Extreme limiting dilution assays (ELDAs) show 2-week pretreatment with CPI-613 100 uM decreases stem cell frequency in GBM cells. Stem cell frequency (1 in n) for each condition shown on plot. 33, 11, 3, 1 cells seeded per well, *n*=10 replicates per condition. Data shown are mean +/− SEM. Data analyzed using Mann-Whitney test **(A and C)**, ELDA Chi-square test **(D)**; ns: not significant, \**P* < 0.05, \*\**P* < 0.01, \*\*\*\**P* < 0.0001.

Previous work has highlighted the importance of mitochondrial function in cancer stem cells [15], however the direct role of mitochondrial respiration in glioma stem cell maintenance is unclear. We show that mitochondrial respiration is perturbed in ALA-treated GBM cells; to further implicate mitochondrial activity in the maintenance of GBM stemness, we employed CPI-613, a mitochondrial respiration inhibitor [16]. As expected, treatment with CPI-613 inhibited overall mitochondrial respiration in a dose-dependent manner (**Supplementary Figure 2C, D**). At 100 μM, cell proliferation is largely unaffected, but mitochondrial respiration is inhibited (**Supplementary Figure 2C, D**); we therefore used this dose for the following ELDA experiments. We pretreated GBM cells with vehicle or with CPI-613 for two weeks and subjected them to ELDA in NSC medium (without CPI-613). ELDA analysis showed that CPI-613 pretreatment (100 μM) attenuates stem cell frequency, recapitulating the stemness reduction seen in ALA-pretreated cells (**Figures 1E and 2D**). These results reveal the critical roles of mitochondrial function in the maintenance of GBM stemness, and suggest compromised mitochondrial function likely contributes to the diminished GBM stemness caused by adenine shortage.

A striking result of the Seahorse XF analyses, shown in Figure 2C, is that ALA treatment eliminates GBM cells’ spare respiratory capacity (also known as reserve respiratory capacity), a measure of cellular fitness [17, 18]. Higher spare respiratory capacity has been associated with radioresistance, resistance to cell death, and lower production of reactive oxygen species (ROS) [17–19]. Furthermore, GBM cells that acquired TMZ resistance were found to display a higher spare respiratory capacity, suggesting the importance of mitochondrial reserve in GBM cells’ response to TMZ [20]. Collectively, these previous findings, together with the weakened stemness and elimination of spare respiratory capacity by ALA, led us to ask whether ALA treatment can sensitize *MTAP*-deficient GBM cells to temozolomide (TMZ).

First, we conducted Seahorse XF analyses treating cells for 24 hours with vehicle, TMZ alone, or TMZ combined with 2-week ALA pretreatment. We observed that with TMZ treatment alone (at 100 μM) for 24 hours, cells maintain a positive reserve capacity (**Figure 3A)**. Furthermore, we saw that ALA-pretreated cells treated with TMZ 100μM maintain the diminished spare respiratory capacity seen in ALA-pretreated-only cells (**Figure 3A**). Interestingly, analysis of extracellular acidification rate (ECAR) showed that 2-week ALA pretreatment, without or with TMZ, reduces cells’ ability to increase ECAR in response to stressed conditions (oligomycin) compared with vehicle or TMZ alone (**Supplementary Figure 2E**). Consistent with these observations, cell energy phenotype analysis shows that ALA-pretreated cells (ALA alone and TMZ + ALA) are unable to shift their metabolism toward an “energetic” phenotype to meet induced energy demands, reflecting diminished cell fitness (**Figure 3B**).

**Figure 3.**
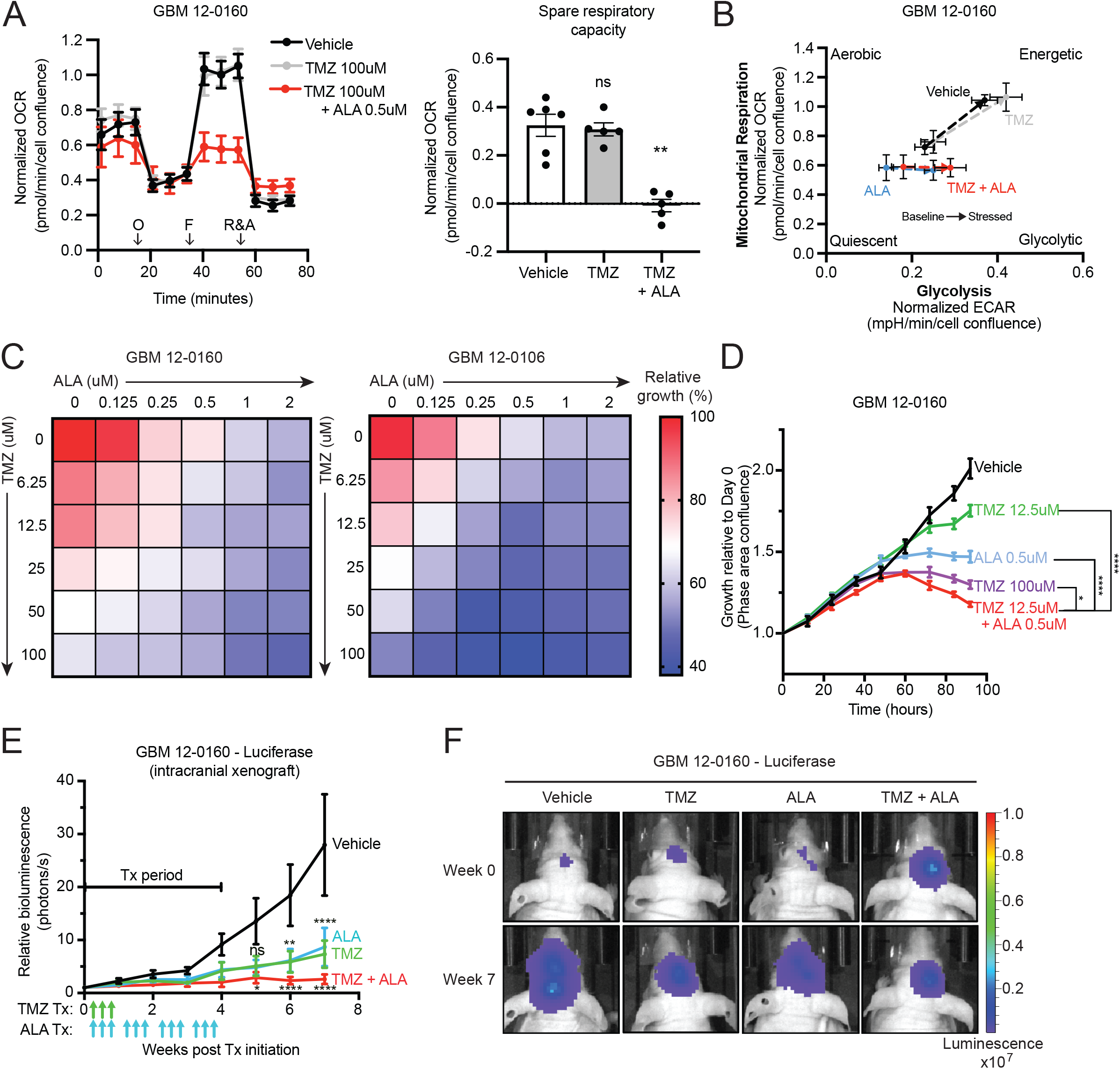
Inhibition of *de novo* adenine synthesis using L-Alanosine (ALA) attenuates *MTAP*-deficient GBM cells’ resistance to temozolomide (TMZ) *in vitro* and *in vivo*. **(A)** Seahorse XF analysis of GBM 12–0160 shows that 2-week ALA 0.5 uM-pretreated cells maintain attenuated maximal respiration (left) and abolished spare respiratory capacity (right) when treated acutely (24 hours) with TMZ. *n*=6 replicates for Vehicle, *n*=5 replicates for each other condition. O: Oligomycin, F: FCCP, R&A: Rotenone and Antimycin A. **(B)** Seahorse XF cell energy phenotype analysis shows that ALA 0.5 uM-pretreated cells (ALA alone and TMZ + ALA) are unable to shift their metabolism toward an “energetic” phenotype to meet induced energy demands. Arrows indicate Baseline to Stressed condition. Stressor compounds: oligomycin (ECAR) and FCCP (OCR). **(C)** Heatmaps show combination effect of TMZ and ALA on growth of two *MTAP*-deficient GBM cell lines. Growth assayed with Incucyte Live Cell Imaging system, 4 days (percentage confluence). Growth shown relative to Vehicle condition, *n*=3 replicates for each drug combination. **(D)** Longitudinal proliferation assay shows low-dose combination TMZ+ALA treatment is more efficacious than single agent treatment and as efficacious as high-dose TMZ alone in GBM-120160. Growth assayed with Incucyte Live Cell Imaging system (percentage confluence). Growth shown relative to time 0 for each treatment condition, *n*=3 replicates. **(E)** Intracranial xenograft of GBM 12–0160 – Luciferase in nude mice given either vehicle, single agent, or combination treatments. Growth curves show combination TMZ and ALA is more efficacious than single agent treatment in inhibiting GBM growth *in vivo*, *n*=5 mice per group. For statistics, each group is compared to Vehicle. *P* value summaries labeled above ALA and TMZ curves (ns, **, ****) represent significance for both ALA and TMZ curves. Tx: Treatment. Green arrows: TMZ administered; Blue arrows: ALA administered. **(F)** Representative bioluminescence images from mice in (E) depicting change in intracranial tumor luciferase signal from Week 0 to Week 7. Data shown are mean +/− SEM. Data analyzed using Mann-Whitney test **(A, right),**Two-Way ANOVA followed by Tukey’s multiple comparisons test **(D-E)**, *P* values labeled in **(D)** represent Tukey’s test comparison for latest time point; ns: not significant, \**P* < 0.05, \*\**P* < 0.01, \*\*\*\**P* < 0.0001.

Next, using paired MTAP-intact and MTAP-deficient cell lines, we established that *MTAP*-deficient GBM cells display increased resistance to TMZ (**Supplementary Figure 3A, B**), agreeing with our previous finding [12]. Subsequently, we subjected two GBM cell lines to various combinations of TMZ and ALA *in vitro*. We used the Incucyte Live Cell Imaging system to track proliferation of cells and assess efficacy of combination treatments. Indeed, we observed that several combinations of TMZ and ALA are more efficacious in inhibiting growth than either TMZ or ALA alone (**Figure 3C**; **Supplementary Figure 4A,**). Importantly, we observed that a low dose (12.5 μM) of TMZ in the presence of a low dose (0.5 μM) of ALA displays efficacy comparable to that of high dose (100 μM) TMZ alone (**Figure 3C, D**). Furthermore, imaging and longitudinal Annexin V apoptosis assay of GBM cells confirm that low-dose combination treatment has similar efficacy to 100 μM TMZ as a single agent (**Supplementary Figure 4C, D**). Collectively, in agreement with the diminished mitochondrial spare respiratory capacity and stemness of GBM cells, these *in vitro* results suggest that ALA can be used to sensitize GBM cells to standard-of-care therapy TMZ.

To further test the effect of TMZ and ALA combination treatment in vivo, we orthotopically implanted *MTAP*-deficient GBM 12-0160 cells expressing luciferase into athymic nude mice, randomized the mice into four groups of treatment: Vehicle (saline), TMZ (5 mg/kg), ALA (150 mg/kg), and TMZ+ALA (TMZ 5 mg/kg + ALA 150 mg/kg), and monitored tumor progression by weekly bioluminescent IVIS imaging (**Supplementary Figure 4E**). We observed that single agent treatment, either TMZ or ALA alone, had moderate efficacy *in vivo* compared to Vehicle (**Figure 3E, F**). Importantly, tumors in the combination treatment group remained stable, with very little growth by week 7 in comparison to those treated with a single agent (**Figure 3E, F**). Collectively, these results suggest that combining TMZ and ALA leads to a more durable response than either TMZ or ALA alone in inhibiting the growth of *MTAP*-deficient GBM *in vitro* and *in vivo*.

Collectively, in this study we exploited the susceptibility of *MTAP*-deficient GBM cells to *de novo* adenine synthesis inhibition and examined the effects of adenine blockade on the maintenance of GBMs’ stemness, mitochondrial function, and response to chemotherapy. Findings from this study advance our understanding of these critical aspects of GBMs in several ways. First, recent work has identified numberous epigenetic and metabolic factors as essential for promoting GBM stemness and consequently as worthwhile therapeutic targets [8–11, 21–23]. In particular, it has been shown that the guanine synthesis arm of purine metabolism can be targeted for overcoming GBM stemness [24]. Our finding that adenine shortage can attenuate the stemness of GBMs illustrates a critical role of the adenine synthesis arm of purine metabolism in promoting GBM stemness and establishes adenine synthesis as an additional therapeutic vulnerability for GBMs. This GBM cell-specific adenine blockade strategy is facilitated by the fact that about half of GBMs bear homozygous *MTAP* deletions, making them deficient in adenine salvage and uniquely susceptible to this therapeutic strategy. Moreover, the suppressive effect of transient adenine synthesis inhibition on GBM stemness further supports the feasibility of this therapeutic strategy.

Second, through transcriptomic and metabolic analyses, we provide direct evidence to support the essential role of mitochondria in maintaining GBM stemness and suggest that attenuated mitochondrial respiration and spare respiratory capacity partially underlie the effects of adenine blockade on stemness. These findings highlight the pathogenic significance of the distinct metabolic profiles in BTICs in contrast to those in non-stem GBM cells [18]. While the transcriptional output from the mtDNA in BTICs is diminished after adenine shortage, we speculate it is more likely this is indicative rather than causative of defective mitochondrial function. The lasting effect of adenine blockade on mitochondrial function in BTICs, as evidenced by measurable defects in mitochondrial function even when the blockade was removed, is particularly intriguing. Further studies illustrating the mechanistic link between adenine supply and mitochondrial function, and/or a metabolic profile-based stratification of GBMs for identifying those most vulnerable to this therapeutic strategy, are needed for further potentiating adenine blockade treatment.

Finally, the therapeutic implication of the aforementioned findings is highlighted by adenine synthesis inhibition-conferred BTIC susceptibility to TMZ, an untargeted, toxic therapy that is currently standard-of-care for GBM patients [25]. While it is possible that the effects of adenine blockade on the DNA damage response contriute to the observed therapeutic benefit [26], we postulate that mitochondrial damage by adenine shortage is part of the mechanism, as the effects of adenine blockade on GBM stemness can be recapitulated by direct pharmacological inhibition of mitochondrial respiration. Adenine blockade serves as another example to highlight the promising principle of exploiting metabolic vulnerabilities for overcoming GBM stemness and resistance to TMZ, as demonstrated by a previous study [27]. We speculate that dissecting the interplay between adenine supply, mitochondria functionality, GBM stemness, and TMZ sensitivity will be critical for exploiting this principle and maximizing therapeutic benefit in GBM treatment.

In summary, our findings suggest that adenine synthesis inhibition can be a strategy to attenuate stemness and TMZ resistance in *MTAP*-deficient GBM cells, and that inhibition of mitochondrial function and elimination of spare respiratory capacity underlie these effects. We propose that the dual benefits of adenine synthesis inhibition – reducing tumor cell stemness and sensitizing them to TMZ – provide strong rationale for further exploiting this approach to treat GBM, particularly as a chemotherapeutic sensitization strategy.

## METHODS

### Cell culture

Patient-derived GBM cells were derived and cultured as previously described [12]. Briefly, GBM 12-0160 and GBM 12-0106 cultures were derived from patient tumor samples with consent (Duke University Brain Tumor Center) and were maintained in human neural stem cell (NSC) medium (STEMCELL, cat# 05751) supplemented with EGF 20 ng/mL, FGF 10 ng/mL, and heparin 2 μg/mL. These cells were plated and passaged on laminin-coated tissue culture plates. Transformation of normal human astrocytes was performed as previously described [12, 28]. Briefly, cells were transduced with four defined genetic factors (*OCT4*, *MYC* (T58A), *HRAS* (G12V), and *TP53* dominant negative) and cultured in NSC medium with 3% FBS. Seven to ten days after transduction, cells were either transduced again with CRISPR/Cas9 lentivirus and sgRNAs targeting *MTAP* (Transformed Astrocyte #6) or treated constitutively with MTAP inhibitor, MTDIA (Transformed Astrocyte #1). Cells were subsequently switched to serum-free NSC medium. All cell lines were maintained in a humidified incubator at 37°C with 5% CO_2_.

### Extreme limiting dilution assay

Sphere forming capacity of GBM cells was assessed using the Extreme Limiting Dilution Assay (ELDA). Decreasing numbers of cells per well (33, 11, 3, and 1, 12 wells per condition) were seeded into 96 well flat-bottom plates in NSC medium with or without ALA. In the pretreatment experiment in Figures 3E-F, GBM suspension tumor spheres were treated with ALA 0.25μM for two weeks in NSC medium in 10cm plates prior to being brought into single cell suspension with Accutase (Innovative Cell Tech, cat# AT-104) and seeded for sphere formation. Two to three weeks after plating, the presence and number of tumor spheres in each well were recorded. ELDA analysis was conducted using the online ELDA software (http://bioinf.wehi.edu.au/software/elda) [14]. Overview of ELDAs is shown in Figure 1D.

### Transcriptomic analysis: mRNA-seq

GBM 12-0160 and GBM 12-0106 cells were treated with Vehicle or ALA 0.25uM for 2 weeks, after which total RNA was extracted using the Quick-RNA Miniprep kit (Zymo Research, cat# R1054). Paired-end 150 bp sequencing was performed by Novogene on an Illumina NovaSeq 6000 machine. Data quality control performed by Novogene; Bowtie 2 was used for alignment and Cufflinks 2.2.2 was used for gene expression profiling. HTSeq and edgeR were used for differential count analysis. For relative FPKM presented in Figure 2A, genes with FPKM values of at least 10 were included.

### mtDNA abundance analysis

GBM 12-0160 and GBM 12-0106 cells were treated with Vehicle or ALA (0.25 uM or 0.5 uM) for 2 days, after which total DNA was extracted using the QIAamp DNA mini kit QIAGEN, cat# 51304). mtDNA-specific primers (*MT-ND3* and *MT-ND4*) were used to amplify mtDNA and *ACTB* primers were used to amplify genomic DNA. Quantitative PCR was performed on a Bio-Rad CFX PCR machine and relative abundance of mtDNA was analyzed.

### *In vitro* proliferation assays

Patient-derived GBM cells were plated in laminin-coated 96 well plates and Transformed Astrocytes were plated in suspension in 96 well plates. All cells were plated in NSC medium with different concentrations of Temozolomide (MedKoo, cat# 100810) and/or L-Alanosine (MedKoo, cat# 200130), or CPI-613 (Cayman Chemical, cat# 16981). Proliferation was assayed by Cell Counting Kit-8 (Dojindo, cat# CK04-20) or by Incucyte Live Cell Imaging system (Essen BioScience) as indicated in figure legends. Apoptosis/Annexin V measurement was performed using the Incucyte Annexin V Green reagent (Essen BioSciences, cat# 4642) in conjunction with the Incucyte Live Cell Imagining system, as recommended by the manufacturer. Length of proliferation assays are indicated in figure legends.

### Seahorse XF analysis

Metabolic flux assays measuring oxygen consumption rate (OCR) and extracellular acidification rate (ECAR) were performed using the Seahorse XFe96 Analyzer (Agilent). Patient-derived GBM cells were seeded at a density of 1 to 1.5 × 10^4^ cells per well in NSC medium in laminin-coated Seahorse XFe96 cell culture microplates. The plate was incubated at 37°C for several hours to allow cells to adhere. Different concentrations of L-Alanosine, Temozolomide, or CPI-613 diluted in NSC medium were added to the wells and incubated for 24 hours at 37°C. The following day, the medium was replaced with Seahorse Phenol Red-free DMEM containing appropriate dilutions of L-Alanosine, Temozolomide, or CPI-613. The plate was incubated in a CO_2_-free 37°C incubator for 1 hour and a Mitochondrial Stress Test (Agilent) was performed according to manufacturer’s instructions. Following the mitochondrial stress test, the plate was imaged in an Incucyte Live Cell Imaging system (Essen BioScience). Cell confluence was quantified for each well and used for normalizing OCR and ECAR measurements.

### *In vivo* drug response studies

Orthotopic intracranial tumors were generated by injecting cells (GBM 12-0160, expressing luciferase) into the right caudate nucleus of female athymic nude mice (Jackson Labs, stock# 007850). Cells were mixed 1:3 with methylcellulose and a total of 1 × 10^5^ cells were implanted into each animal. Tumors were allowed to form for 4 weeks after implantation. To establish baseline tumor bioluminescence, mice were injected with D-luciferin 15 mg/kg (GoldBio, cat# LUCNA) and scanned in an IVIS Lumina XR imager. Mice were then randomized into four groups and drug treatment initiated. Mice received three intraperitoneal (IP) injections of TMZ 5 mg/kg (MedKoo, cat# 100810) during the first week of treatment followed by three weeks off, representing one cycle of TMZ treatment. Mice received three IP injections of ALA 150 mg/kg (MedKoo, cat# 200130) weekly for the duration of the treatment period. Saline IP injections were used as the vehicle control treatment. *In vivo* drug response was monitored by weekly bioluminescent imaging of mice as described above and analyzed using Living Image software.

### Synergy/antagonism analysis

To determine possible synergistic or antagonistic effects of TMZ and ALA combination treatment, proliferation data from the Incucyte Live Cell Imaging system were analyzed using Combenefit software (www.cruk.cam.ac.uk/research208groups/jodrell-group/combenefit) [29]. The synergy/antagonism score for each combination was calculated by Combenefit using the Loewe synergy model, where a score >0 indicates synergy, a score of 0 indicates additive effects, and a score <0 indicates antagonism.

### Statistics

All *in vitro* experiments in this study were performed independently at least twice. Data presented as mean +/− standard error of the mean (SEM). Mean values between two groups were compared using t-tests (two-tailed, unpaired, with Welch’s correction) when data were assumed to follow a normal distribution, otherwise non-parametric Mann-Whitney test was performed. Multiple groups were compared using ANOVA, followed by Tukey’s or Sidak’s multiple comparisons tests. Statistical test results were deemed significant if *P* < 0.05. With the exception of ELDA experiments and Combenefit synergy analysis, all statistical analyses were calculated using Graphpad Prism 8 software.

### Study approval

All animal experiments were performed in accordance with protocols approved by the Duke University Institutional Animal Care and Use Committee (IACUC).

## AUTHOR CONTRIBUTIONS

S.X.S. and Y.H. conceived and designed this study. S.X.S. and Y.H. developed the methodology. S.X.S., R.Y., K.R., L.J.H., C.D., P.K.G., and C.J.P. contributed to the acquisition of data. L.H.C. processed and analyzed mRNA-seq data. S.X.S. and Y.H. analyzed and interpreted the data. S.X.S. wrote the manuscript. S.X.S., C.J.P., and Y.H. edited the manuscript with input from all authors. P.K.G. and Y.H. contributed to administrative, technical, or material support. Y.H. supervised this study.

## ACKNOWLEDGMENTS

We thank the Duke Cancer Center Isolation Facility (CCIF/DCI) for their animal maintenance support in this study. We thank Amanda Nichols and Dr. Nancie MacIver at the Duke Cellular Metabolism Analysis core facility for their help performing Seahorse XF analyses. Vector images (round cell culture plate, multi-well plate, clipboard checklist, mouse, camera) used in Figure 1D and Supplementary Figure 4E were designed by Freepik, Dimitry Miroliubov, and Good Ware on flaticon.com. This work was supported by the Department of Pathology and the Preston Robert Tisch Brain Tumor Center at Duke University, as well as by the National Institute of Neurological Disorders and Stroke of the National Institutes of Health, Award Number R01NS101074 (YH).

---

**Supplementary Figure 1.**
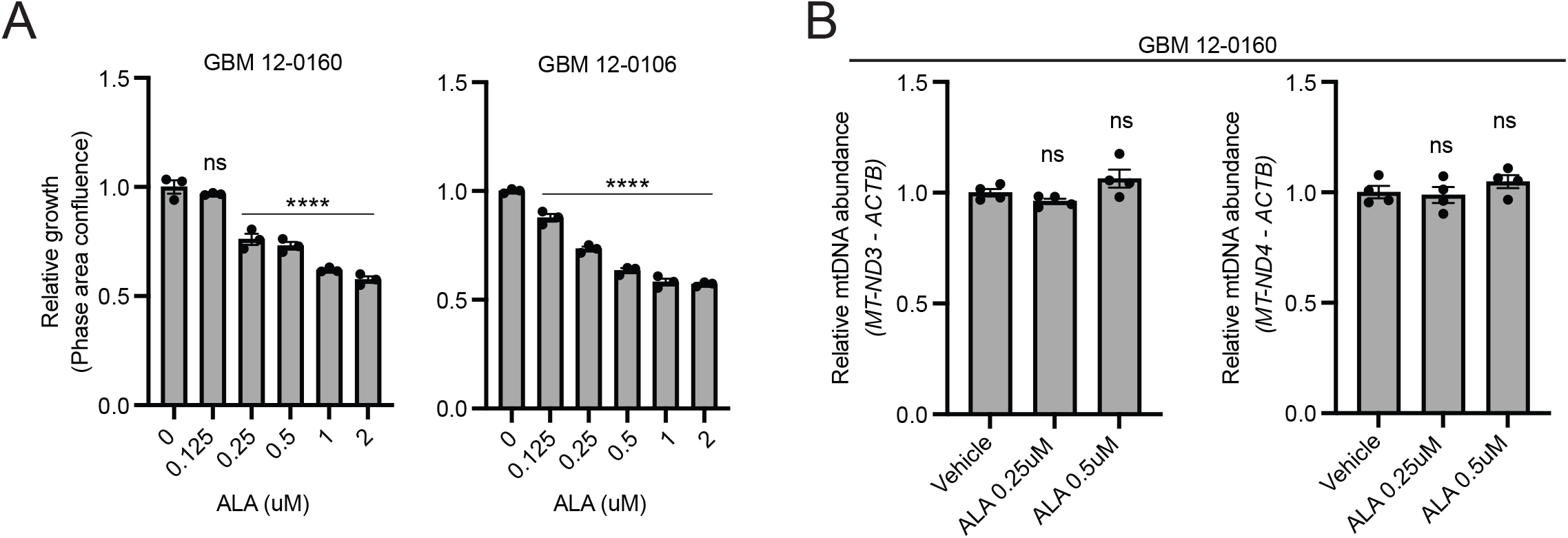
Effects of L-Alanosine (ALA) treatment on proliferation and mtDNA abundance in GBM cells. **(A)** Drug sensitivity plots for ALA in two GBM cell lines. Growth assayed with Incucyte Live Cell Imaging system, 4 days (confluence relative to Day 0, normalized to ALA 0 uM), *n*=3 replicates for each dose. **(B)** Quantitative PCR analysis shows that mtDNA abundance is unaffected by ALA treatment (2 days). *MT-ND3* or *MT-ND4* amplification relative to *ACTB*, *n*=4 replicates per condition. Data shown are mean +/− SEM. Data analyzed using One-way ANOVA followed by Holm-Sidak’s multiple comparisons test **(A)**, *P* values represent comparison of each dose to vehicle (zero); Mann-Whitney test **(B)**; ns: not significant, \*\*\*\**P* < 0.0001.

**Supplementary Figure 2.**
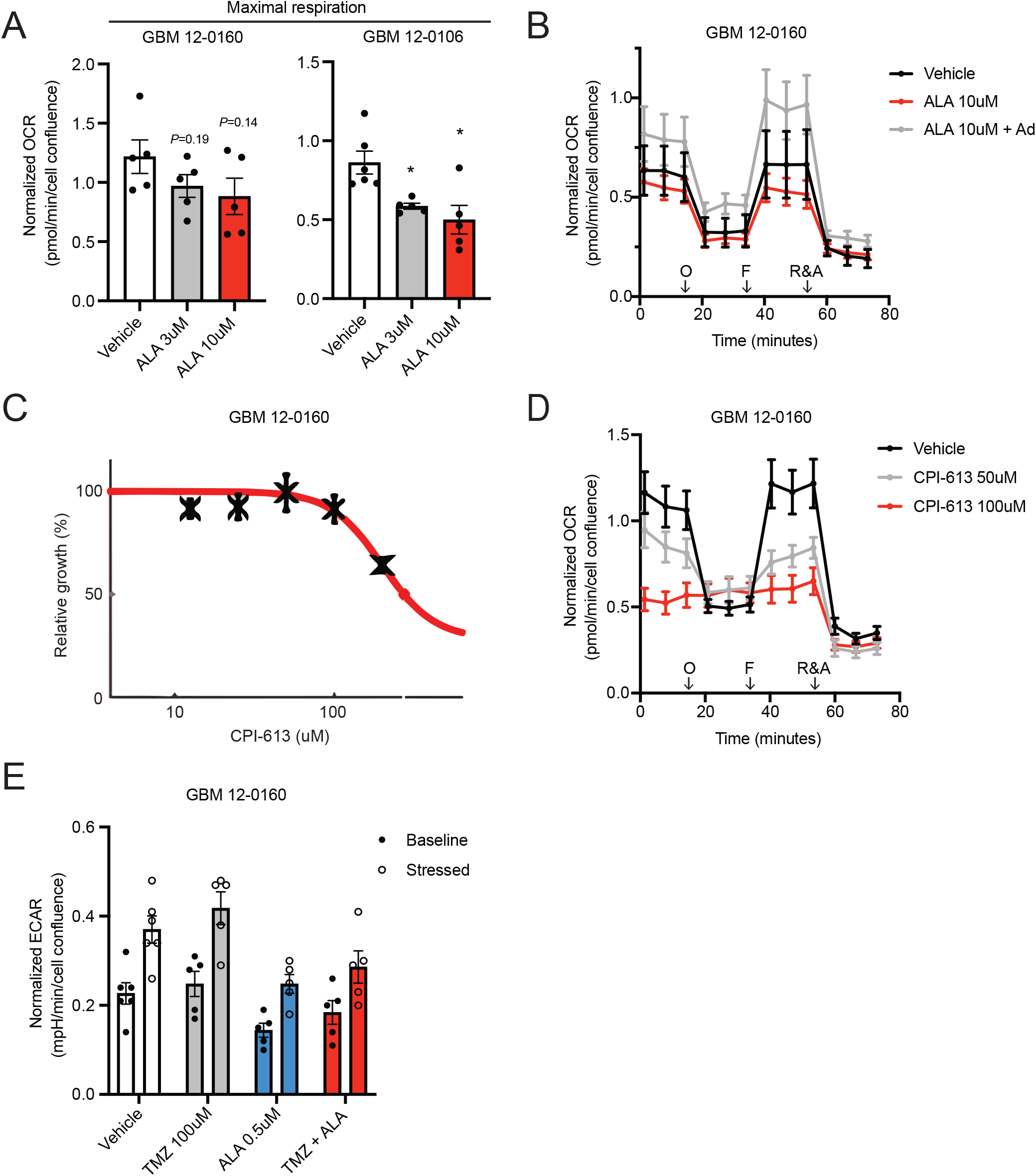
Seahorse XF analyses of L-Alanosine (ALA)- or CPI-613-treated GBM cells. **(A)** Acute (24 hours) ALA treatment reduces maximal respiration in GBM cells. *n*=6 replicates for Vehicle (GBM 12–0106), *n*=5 replicates for each other condition. **(B)** Reduced maximal respiration in acutely (24 hours) ALA-treated cells can be rescued by Adenine (Ad) 10 uM. *n*=6 replicates for Vehicle, *n*=5 replicates for each other condition. O: Oligomycin, F: FCCP, R&A: Rotenone and Antimycin A. **(C)** CPI-613 dose response curve. Plot generated by Combenefit software, mean shown of *n*=3 replicates. **(D)** Acute (24 hour) CPI-613 treatment reduces basal and maximal respiration. *n*=6 replicates for Vehicle, *n*=5 replicates for each other condition. O: Oligomycin, F: FCCP, R&A: Rotenone and Antimycin A. **(E)** Two-week ALA 0.5uM pretreatment reduces cells’ ability to increase ECAR in response to stressed conditions (oligomycin). *n*=6 replicates for Vehicle, *n*=5 replicates for each other condition. Data shown are mean +/− SEM. Data analyzed using Mann-Whitney test **(A)**; \**P* < 0.05.

**Supplementary Figure 3.**
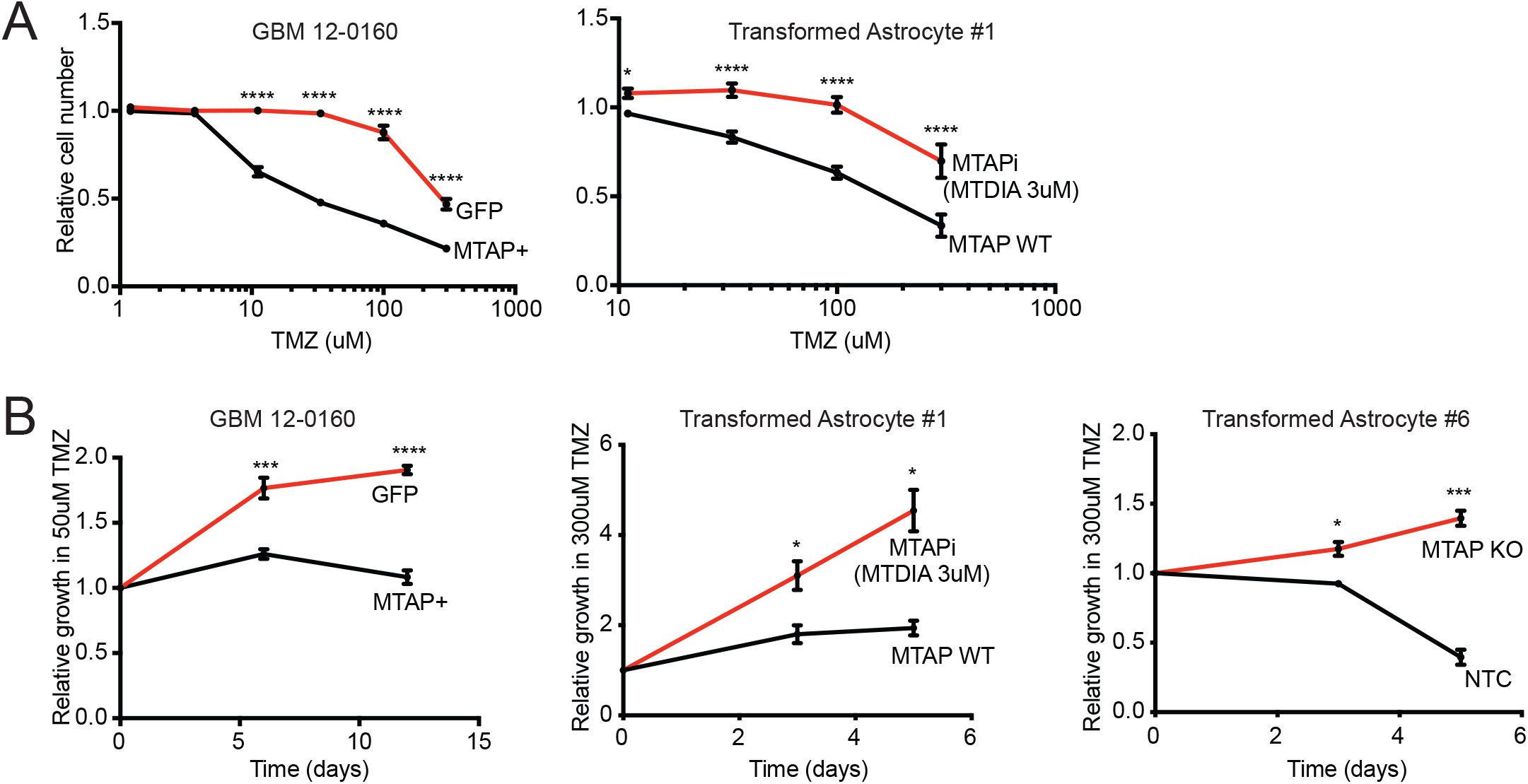
MTAP loss facilitates temozolomide (TMZ) resistance. **(A)** Drug sensitivity curves show GBM 12–0160 (left) and Transformed Astrocyte #1 (right) cells are more resistant to increasing doses of TMZ when MTAP is deficient (left) or pharmacologically inhibited (right). CCK8 assay, 4–6 days, *n*=4 replicates. **(B)** Proliferation assays show genetic absence (left, right) or pharmacological inhibition (middle) of MTAP facilitate TMZ resistance in several GBM models. Manual cell counting using hemocytometer, *n*=3 replicates. Data shown are mean +/− SEM. Data analyzed using Two-Way ANOVA followed by Sidak’s multiple comparisons test **(A)** or multiple unpaired t-tests **(B)**; \**P* < 0.05, \*\*\**P* < 0.001, \*\*\*\**P* < 0.0001.

**Supplementary Figure 4.**
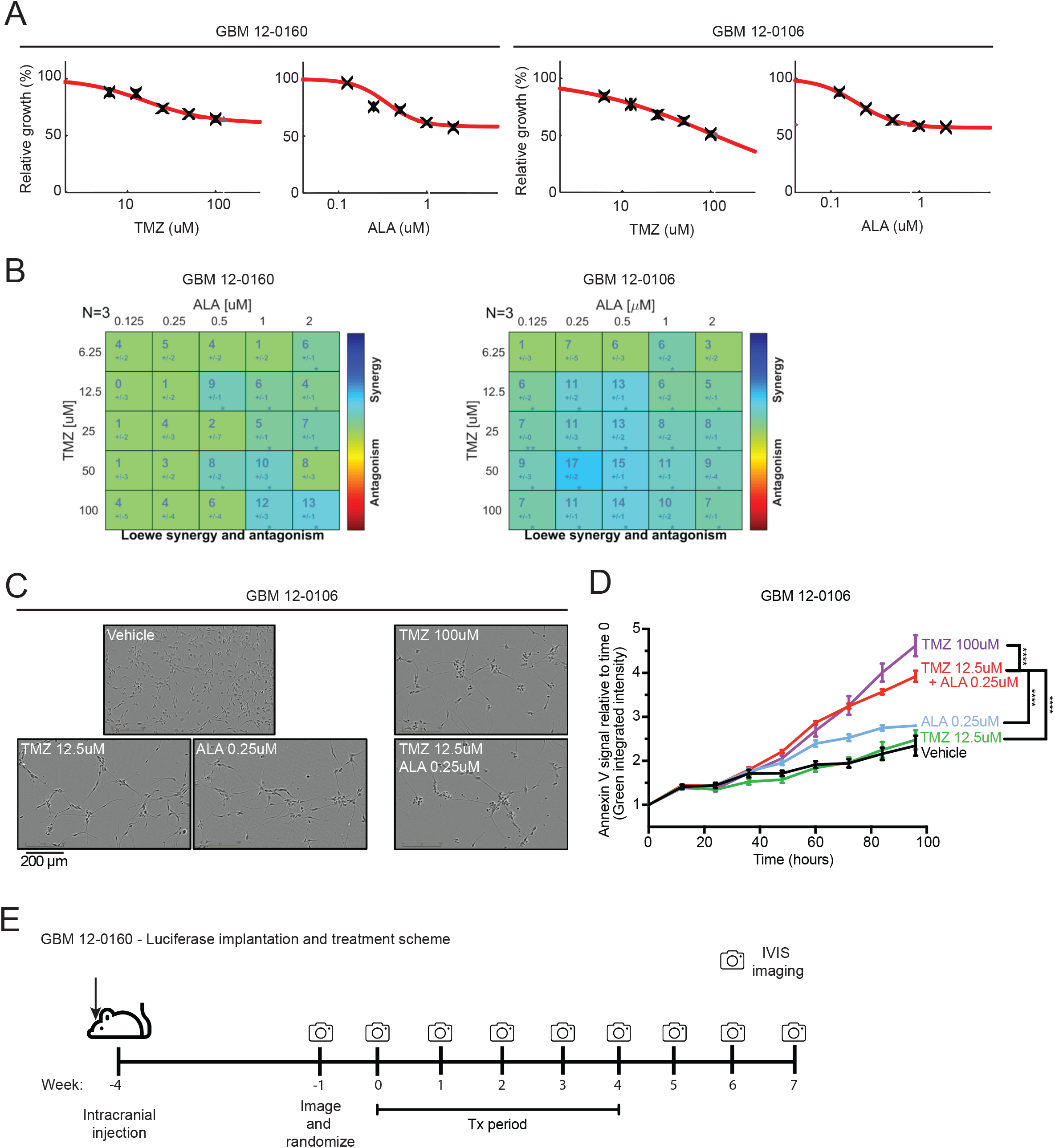
Synergy/antagonism and apoptosis analyses of temozolomide (TMZ) + L-Alanosine (ALA) combination treatment. **(A)** Single agent TMZ and ALA dose response curves for GBM cells. Plots generated by Combenefit software, mean shown of *n*=3 replicates. **(B)** Synergy/antagonism analysis of TMZ and ALA combination treatment data from Figure 3C. Each square indicates the synergy/antagonism score +/− SEM for a given dose combination after 4 days of treatment. Matrix plots generated by Combenefit software, *n*=3 replicates for each drug combination. **(C)** Representative images of adherent GBM 12–0106. Images taken with Incucyte Live Cell Imaging system. Scale bar, 200μm. **(D)** Longitudinal Annexin V staining assay shows low-dose combination TMZ+ALA treatment leads to higher Annexin V signal than single agent treatment and similar signal as high-dose TMZ alone. **(E)** Scheme for *in vivo* drug response experiment using GBM 12–0160 – Luciferase. Data shown are mean +/− SEM. Data analyzed using Combenefit software **(B)**, Two-Way ANOVA followed by Tukey’s multiple comparisons test **(D)**; \**P* < 0.05, \*\**P* < 0.01, \*\*\*\**P* < 0.0001.

